# Coevolutionary dynamics of viruses and their defective interfering particles

**DOI:** 10.1101/2025.10.22.683971

**Authors:** Shiv Muthupandiyan, John Yin

## Abstract

Defective interfering particles (DIPs) are viral mutants that arise naturally during infection. Because they lack one or more essential functions, DIPs cannot replicate on their own, but they can parasitize intact viruses during co-infection by competing for growth resources, thereby interfering with viral replication. The evolutionary interplay between viruses and their DIPs involves growth, mutation, interference, and resource trade-offs, but the mechanisms shaping population-level outcomes remain poorly understood. To address this, we developed a continuous phenotype-space model using coupled partial differential equations that incorporate mutation, phenotype-dependent interference, intrinsic fitness costs, and *de novo* DIP generation. Unlike traditional strong-selection models, this framework captures strong-mutation regimes in which both virus and DIP populations diffuse through trait space and interact based on phenotypic similarity. Our analysis reveals two levels of dynamics. At the population level, viruses and DIPs undergo oscillations, consistent with predator–prey–like cycles (the von Magnus effect) observed experimentally. At the trait level, evolution drives shifts in resistance and interference, producing coevolutionary chases in which viruses temporarily escape and new DIPs emerge to follow, as observed in serial-passage evolution studies. Systematic variation of parameters reveals four qualitative regimes: viral–DIP coexistence, sustained coevolutionary (Red Queen) chase dynamics, DIP extinction, and mutual extinction. Chase dynamics are most strongly promoted by intermediate interference strength and low decay rates, while higher levels drive collapse of one or both populations. The model further predicts thresholds where viral escape is either constrained by intrinsic fitness penalties or enabled through phenotypic separation from DIPs. These findings establish a general framework for virus–DIP coevolution, showing how both population dynamics and trait evolution shape outcomes, with implications for designing DIP-based therapeutics that better resist viral escape.

**Author summary:** Most viruses produce defective versions of themselves, known as defective interfering particles (DIPs). DIPs cannot multiply on their own, but when they infect the same cell as an intact virus, they steal resources and limit the virus’s growth. This makes them a promising antiviral therapy. But viruses may evolve to reduce the ability of DIPs to steal resources. We built a mathematical model that lets both viruses and DIPs vary in traits—such as how quickly they grow or how strongly they interfere—and evolve such traits over time. Our results reveal two layers of dynamics. At the population level, viruses and DIPs can rise and fall in repeating cycles that resemble predator–prey interactions, even when traits stay fixed. With evolution, however, the contest gains another dimension: viruses may shift traits to escape interference, while new DIPs can arise and adapt to follow them. This co-evolutionary chase is a novel feature that sets DIPs apart from conventional therapies, which cannot adjust to viral change. By exploring many conditions, we identified when viruses and DIPs coexist, when one eliminates the other, and when they remain locked in long-term pursuit. These insights suggest principles for designing therapeutic DIPs that might better resist viral escape.

## Introduction

Since the earliest days of virus cultivation, laboratory virus stocks contained aberrant particles that suppressed the replication of fully infectious virus. In a landmark series of experiments, Preben von Magnus showed that undiluted passage cultures of influenza virus led to the accumulation of non-infectious “incomplete” particles that interfered with standard virus replication [1–3]. This progressive enrichment of defective particles and corresponding drop in infectious virus output came to be known as the von Magnus effect [4]. Studies of vesicular stomatitis virus (VSV) and influenza virus showed how the von Magnus effect could drive predator–prey–like oscillatory dynamics between populations of standard virus and defective interfering particles [5–10].

Defective interfering particles, also called DI particles or DIPs, package and deliver defective virus genomes to host cells. The defective viral genomes are byproducts of error-prone genome replication by most RNA viruses, including VSV, influenza, and coronavirus, where their defects are most often linked to the deletion of one or more essential genes for virus growth [11–13]. Sequencing and analyses of environmental and clinical isolates has found defective genomes for many other RNA viruses, providing evidence for their occurrence and transmission in nature as well as their potential to modulate disease severity [14–18].

Early recognition of DIPs as natural parasites of virus replication spurred efforts to understand and exploit their interference for therapeutic benefit. In animal models, DIPs protected mice and ferrets challenged with RNA viruses such as VSV, Semliki Forest virus (SFV), and influenza A virus [19–21]. More recently, DIP-inspired therapeutic interfering particles derived from influenza, polio, SARS-CoV-2, and HIV-1 have shown promise in rodent and non-human primates [22–25]. These therapeutic interfering particles (TIPs) may suppress the emergence of viral resistance through potent interference [24, 26], selection pressures that limit escape [27], or activation of broadly protective host immunity [23]. In nature, DIPs arise as byproducts of error-prone viral replication, and the same mutational processes that generate viral diversity also drive diversity among DIPs. A key unanswered question is how ecological and coevolutionary interactions between viruses and their DIPs shape viral growth, disease severity, and viral persistence [28].

The potential evolution of viral resistance poses a significant challenge to DIP-based therapies. Early experimental work with VSV demonstrated that viruses can indeed evolve to escape DIP interference [29]. Time-shift experiments, where DIPs were tested against viruses from past, present, and future passages [30], revealed a dynamic of transient resistance [31]. The virus was resistant to contemporary DIPs but remained susceptible to those from the future or distant past. The specific mechanism driving this resistance was later identified as mutations in the viral polymerase that altered its binding specificity, rendering existing DIPs less effective [32].

Strategies to prevent this viral escape have been explored theoretically. A key concept is an “evolutionary conflict,” where competing selection pressures make resistance a self-defeating strategy for the virus. For instance, within-host models for HIV show that any viral mutation that evades a TIP by reducing the production of a shared protein, such as the viral capsid that packages the viral genome, would simultaneously impair the virus’s own replication fitness, creating a strong selective pressure against escape [33]. This principle was later shown to be robust at the population level, where additional evolutionary trade-offs further constrain the spread of resistant mutants [27]. A critical assumption underlying these frameworks, however, is that viral escape occurs through specific, discrete mutations against a static therapeutic agent. Therefore, investigating a “strong-mutation” environment, where a continuous flux of mutations allows for the evolution of both the virus and its interfering particles themselves, requires a different mathematical approach.

To investigate coevolutionary interactions between viruses and their DIPs, we build on recent advances in evolutionary modeling. Classical adaptive dynamics assume rare mutations and monomorphic populations interrupted by discrete trait substitutions, but these assumptions may not be valid in virus–DIP systems where mutations occur frequently and generate broad polymorphism within a single infection cycle [34]. For such strong-mutation regimes, new theoretical work develops macroscopic models that treat mutation as a diffusion process in phenotype space, allowing continuous adaptation of entire trait distributions rather than stepwise shifts [35, 36]. This perspective is essential here because interference depends on phenotypic similarity, and the evolutionary outcome hinges on the joint movement of virus and DIP distributions, not on isolated genotypes.

Within this context, the Red Queen hypothesis—named after Lewis Carroll’s character who declares, “it takes all the running you can do to stay in the same place” —emphasizes that coevolution is driven by reciprocal rather than externally imposed selection [37]. In this view, antagonistic species must continuously adapt just to maintain their relative fitness. One manifestation, the “Chase Red Queen,” describes directional pursuit through multidimensional phenotype space: one population evolves to reduce its exploitable similarity to the other, while its antagonist evolves to close the phenotypic gap, leading to a persistent chase rather than oscillation around a fixed optimum [38]. Recent theoretical work has extended this idea to host–pathogen systems by modeling phenotype-structured populations and demonstrating how such chases can arise intrinsically from coevolutionary feedback. [39].

Building on these insights, we modify mathematical modeling of host-pathogen systems to explore virus–DIP coadaptation under strong mutation. We hypothesize that their dynamics may exhibit a Chase Red Queen pattern, with parameters such as mutation rate and interference strength determining whether the outcome is viral containment, escape, or extinction. By charting these coevolutionary regimes, we provide a theoretical basis for engineering TIPs that leverage trade-offs to maintain long-term viral control.

## Methods

We model DIP-virus coevolution in strong-mutation environments where interference strength varies with phenotypic distance between DIPs and viruses. Building on the framework of Alfaro et al. [39], we incorporate interactions across phenotype space and *de novo* DIP generation, capturing coevolutionary dynamics between phenotypically distant populations.

The model tracks the population densities of the virus, *v*(**x**, *t*), and DIP, *d*(**x**, *t*), as they evolve over a two-dimensional phenotype space, **x** ∈ ℝ^2^. We define this space abstractly to represent any set of continuous traits that could affect interference, rather than tying them to a specific biological mechanism. Examples could include affinity of virus or DIP genomic templates for the viral replicase or packaging proteins, or their replication rates (Fig 1). The evolution of these densities is driven by their phenotypic interactions, namely interference and competition for shared resources.

**Fig 1.**
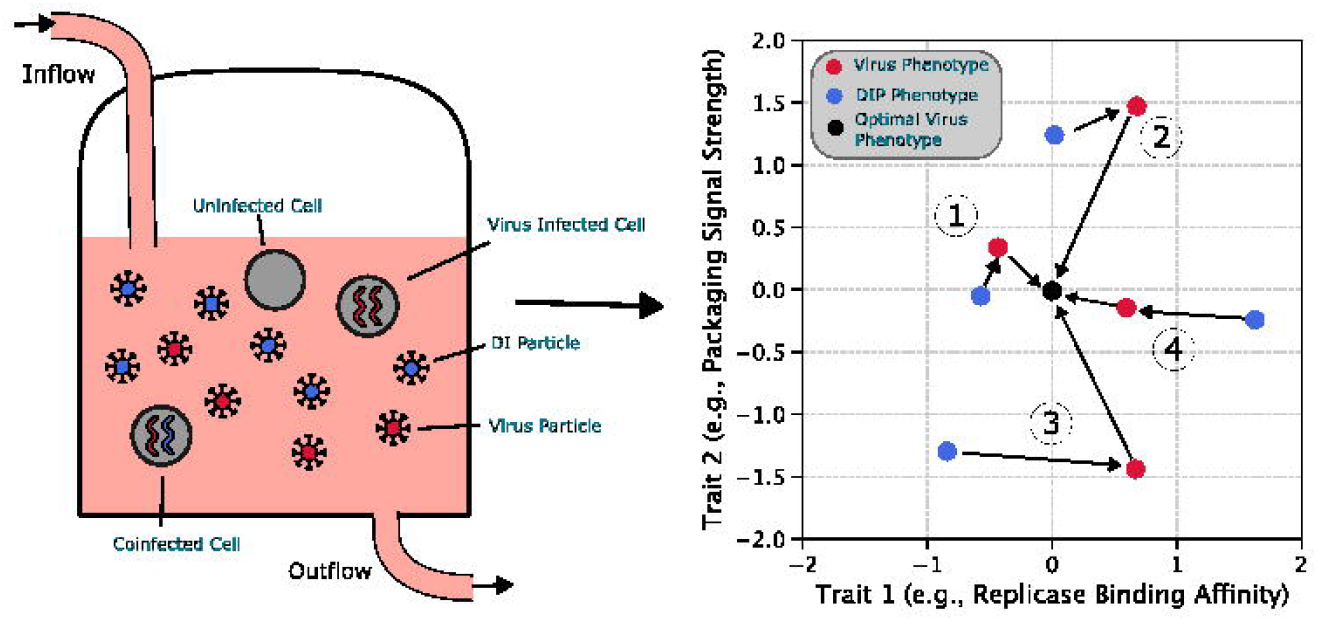
Schematic representation of the virus-DIP coevolutionary system. **Left panel:** Experimental setup modeled as a well-mixed bioreactor with continuous inflow of fresh cells and media, and outflow removing dead cells, viruses, and DIPs. Viral genomes (red) and defective viral genomes (blue) are packaged into particles that compete for cellular resources under finite carrying capacity constraints. **Right panel:** Abstract two-dimensional phenotype space used in the mathematical model, where each point represents a distinct viral (red) or DIP (blue) phenotype characterized by continuous traits (e.g., replicase binding affinity, packaging signal strength). The origin represents the fitness optimum for viruses, while proximity between virus (red) and DIP (blue) populations determines DIP fitness. Numbered scenarios represent: (1) virus retains high intrinsic fitness at the optimum but suffers strong interference from a co-localized DIP population; (2) virus has moved far from the fitness optimum but experiences strong DIP interference due to close phenotypic proximity; (3) virus has low intrinsic fitness far from the optimum and DIP has limited replication capacity due to weak interference; (4) virus maintains high intrinsic fitness near the optimum while DIP interference remains weak due to phenotypic distance.

We model interference based on the principle of phenotypic similarity: the closer DIPs are to the virus in phenotype space, the more strongly they interfere. This principle can be captured mathematically using convolution integrals, which represent weighted averages of interactions across a neighborhood. In spatial population dynamics this formalism was developed to describe how organisms interact more strongly with nearby individuals than with distant ones [40]. Here, we extend the same idea into phenotypic space, where “distance” reflects trait differences rather than physical separation. The biological motivation for this principle is strong. For example, phenotypic divergence is known to reduce interference in VSV, where mutations to the viral replicase can abolish its ability to bind to DIP templates [32]. Likewise, the efficacy of DIP-mediated “genome stealing” in HIV-1 is determined by the similarity of their dimerization initiation sequences [33]. Our model abstracts this concept by using a Gaussian kernel, *G*(**x**), to represent the interaction strength as a function of Euclidean distance in phenotype space. The total interference cost to the virus, *I*_*V*_, and the corresponding benefit to the DIP, *I*_*D*_, in a well-mixed system are given by:

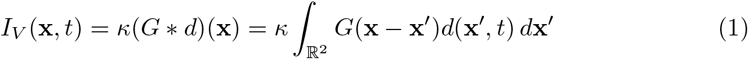

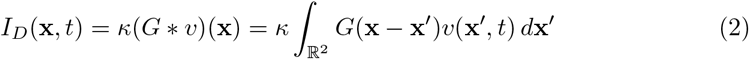

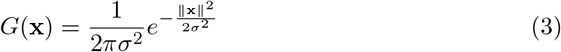

where *κ* is an interference strength constant, and the kernel *G*(**x**) (3) has a scale, *σ*, that controls how rapidly interference declines with phenotypic distance. All variables and parameters are summarized in Table 1 and Table 2.

**Table 1.** Dimensional variables, parameters, and functions.

| Symbol | Units | Description |
| --- | --- | --- |
| **State variables** |  |  |
| $v(\mathbf{x}, t)$ | particles (phen. unit)<sup>-2</sup> | Virus density in phenotype space |
| $d(\mathbf{x}, t)$ | particles (phen. unit)<sup>-2</sup> | DIP density in phenotype space |
| $V(t)$ | particles | Total virus population |
| $D(t)$ | particles | Total DIP population |
| $\bar{\mathbf{x}}_V(t)$ | phen. unit | Mean virus phenotype (distribution centroid) |
| **Functions / operators** |  |  |
| $G(\mathbf{x})$ | (phen. unit)<sup>-2</sup> | Gaussian kernel |
| $\nabla^2$ | (phen. unit)<sup>-2</sup> | Laplacian operator for mutation (diffusion) |
| **Parameters** |  |  |
| $r_V$ | time<sup>-1</sup> | Intrinsic viral replication rate |
| $\mu$ | (phen. unit)<sup>2</sup> time<sup>-1</sup> | Mutation rate (phenotypic diffusion) |
| $\alpha$ | time<sup>-1</sup> (phen. unit)<sup>-2</sup> | Strength of selection toward the origin |
| $\beta$ | time<sup>-1</sup> (phen. unit)<sup>-2</sup> | Strength of the aggregation penalty |
| $\eta$ | time<sup>-1</sup> | Rate of *de novo* DIP generation |
| $\kappa$ | (phen. unit)<sup>2</sup> particles<sup>-1</sup> time<sup>-1</sup> | Per-particle interference strength |
| $\gamma$ | time<sup>-1</sup> | Background removal (decay + washout) rate |
| $\sigma$ | phen. unit | Interaction width of the Gaussian kernel |
| $K$ | particles | Carrying capacity (total particles) |

**Table 2.**
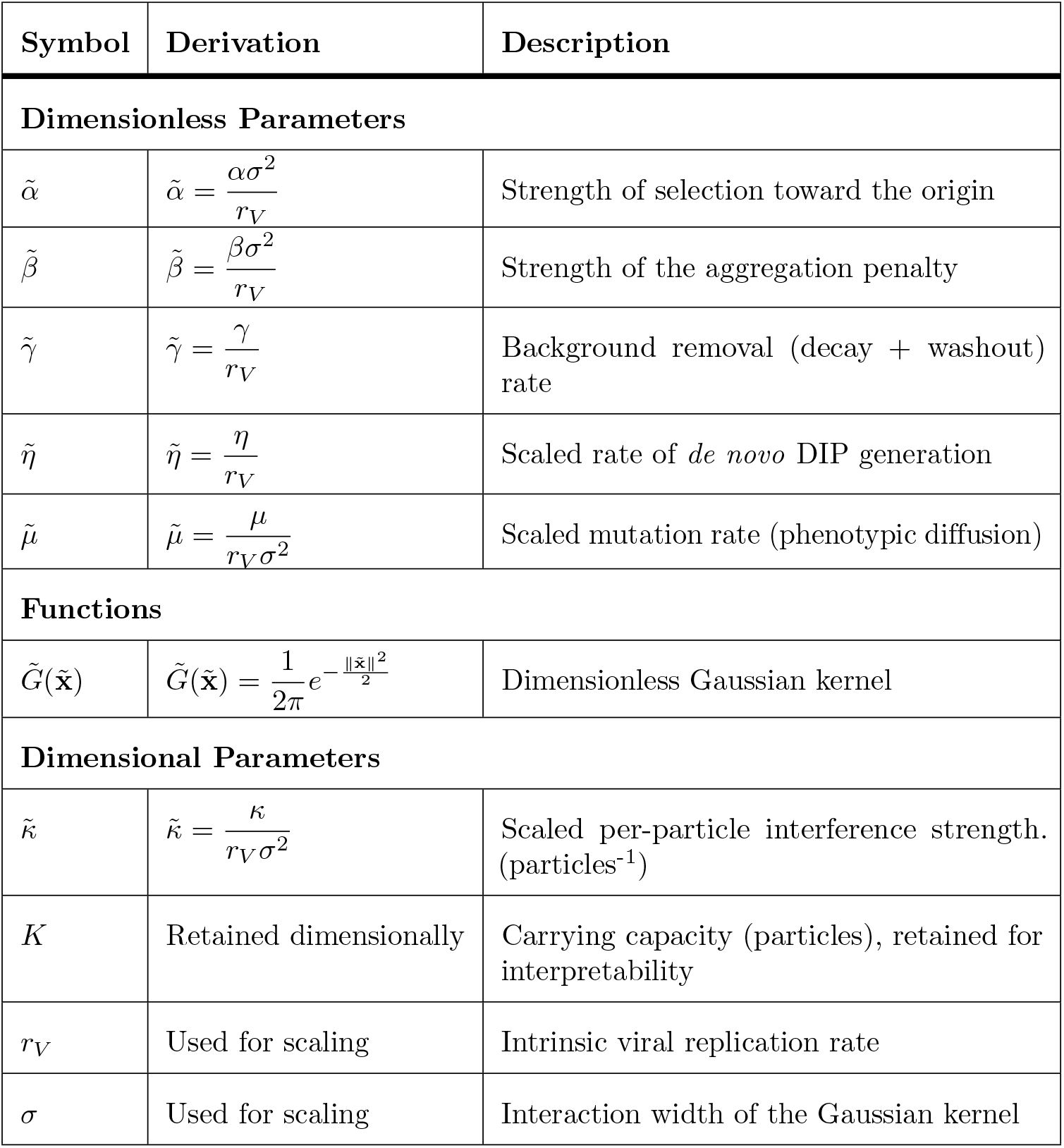
Nondimensionalized model parameters.

The interference terms are designed to represent a zero-sum resource transfer. The cost incurred by the virus population, Eq (1), directly corresponds to a gain for the DIP population, Eq (2), mirroring the direct hijacking of viral replication machinery. This conservation holds at the population level, not necessarily at a specific phenotype. The total resources lost by the virus population are equal to the total resources gained by the DIP population, as shown by the integral equality:

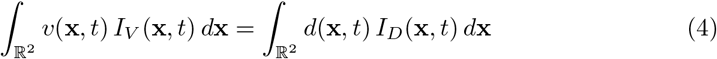

This identity is a direct consequence of the symmetric structure of the convolution integrals. By substituting Eq (1) and Eq (2) and swapping the order of integration, both sides of the equation simplify to the same expression:

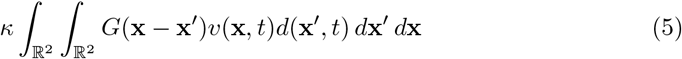

We selected this zero-sum case as a neutral baseline because the net efficiency of resource hijacking is ambiguous. For instance, a DIP could be more efficient than a virus if its smaller genome enhances replication speed (a “positive-sum” outcome) [41]. Conversely, it could be less efficient if its deletions interfere with key functions like viral encapsidation or DIP self-interference (a “negative-sum” outcome) [42, 43]. Therefore, a one-to-one transfer provides the clearest foundation for isolating the dynamics of competitive interference.

To account for finite resources, we regulate overall population growth using a logistic term with a carrying capacity, *K*. This framework is well aligned with microbial culture systems in which a fixed-volume bioreactor maintains a constant carrying capacity as fresh cells continuously flow in [44, 45] (Fig 1). Serial passaging represents its discrete, batch-like counterpart, in which a finite number of host cells limits viral expansion in each cycle. For virus and DIP populations, *V* and *D*, the logistic factor, *L*, is given by:

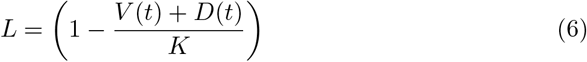

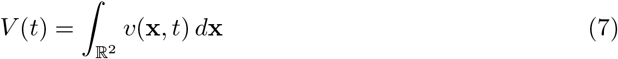

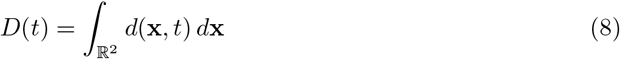

For simplicity, this formulation assumes that virus and DI particles contribute equally to saturating *K*, though in reality, DIPs harboring shorter genomes may be less resource-intensive per particle.

We model the virus’s intrinsic fitness landscape by adapting the framework of Alfaro et al. [39]. This landscape is composed of two quadratic penalty terms. The first, − *α* ∥**x** ∥^2^, establishes a fixed fitness optimum at the origin, penalizing viruses for deviating from this global peak. The second is an aggregation term, 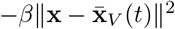, which penalizes viruses for straying from the population’s current mean phenotype, 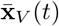This mean phenotype is calculated as:

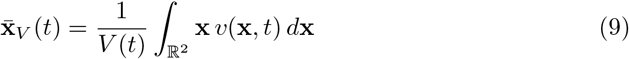

Mathematically, the aggregation term ensures population cohesion and prevents non-biological ring structures and fragmentation [39]. Viral populations are understood to function as cohesive “quasispecies,” existing not as a single genotype but a cloud of related mutants centered around a dominant “master copy” [46]. The ensemble provides complementation benefits that individual clones lack [47]. A virus that becomes phenotypically isolated from this swarm loses these benefits, incurring a fitness cost. Therefore, the *β* term models a plausible selective pressure that penalizes phenotypic outliers, promoting the cohesion that may be necessary for the quasispecies to act as a collective unit of selection.

We also account for the *de novo* generation of DIPs from replication errors in the virus population. These errors produce a spectrum of outcomes. Some errors produce viable virus variants whose small phenotypic shifts are captured by our diffusion term (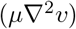). Other errors can be lethal, creating non-viable particles that cannot replicate at all and are thus excluded from our model. To model the *de novo* generation of DIPs we focus on the critical intermediate class of replication errors: defective genomes that are replication-incompetent on their own but retain the necessary sequences to be replicated and packaged by the machinery of the standard virus. This generation is modeled as a transfer of density from the virus to the DIP population at a rate *η*. A key assumption of this term is that a newly generated DIP inherits the exact phenotype of its parent virus. We ground this assumption in the requirement that for a DIP to be conditionally replication-competent, it must retain significant functional similarity to its parent. Major phenotypic changes would likely render an emergent DIP non-viable and thus irrelevant to the coevolutionary dynamic. This assumption therefore focuses the model on the subset of *de novo* DIPs that are most likely to interfere.

Mutation is modeled as diffusion in phenotype space, analogous to the process of diffusion of particles in a physical space. The random motion is caused by the random changes to phenotype by genetic mutations. An initially clonal population will therefore “diffuse” or spread out across the phenotype space over time. We mathematically represent this with the Laplacian operator, 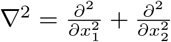 [36]. For simplicity, we assume this diffusion is isotropic and that both the virus and DIP populations share the same diffusion coefficient, the mutation rate parameter, *µ*.

We define the intrinsic viral growth rate as *r*_*V*_. We then assume particles can be removed from the population in two ways. The first is intrinsic biological decay, such as the natural degradation or inactivation of viral particles [48]. The second is physical removal from the system, corresponding to the continuous dilution in experimental setups like the constant-volume bioreactor (Fig 1). In such a system, this physical washout rate is defined as the outflow rate divided by the constant fluid volume of the bioreactor. For simplicity, our model combines these biological and physical removal mechanisms into a single constant rate, *γ*, applied equally to both virus and DIP populations.

The dynamics of virus density, *v*(**x**, *t*), and DIP density, *d*(**x**, *t*), are described by the following system of coupled partial differential equations, which incorporates all the components described above and summarized in Table 1.

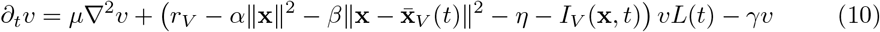

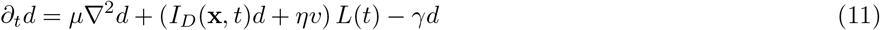

To reveal the fundamental dynamics of the system and reduce the number of free parameters, we non-dimensionalize time and phenotypic space. We introduce the following dimensionless quantities, denoted with a tilde:

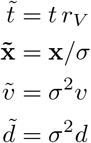

This transformation recasts the model such that time is measured in units of the viral replication cycle and phenotypic distance is measured in units of the interaction width *σ* (i.e., in standard deviations of the Gaussian kernel). To preserve direct interpretability of virus and DIP population sizes and to enable comparison to experimental particle counts, we retain the carrying capacity *K* as a particle count.

Thus, substituting the definitions from Table 2 into the original model (Eqs (10) and (11)), and then dropping the tildes for clarity, yields the system we examine for the remainder of the paper:

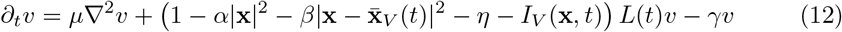

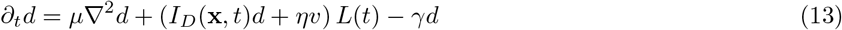

We formally state the initial-boundary value problem that forms the basis of our analysis. The goal is to find the non-negative population densities *v*(**x**, *t*) and *d*(**x**, *t*) satisfying Eqs (12) and (13). For our numerical implementation, this system is solved on a bounded rectangular domain Ω ⊂ ℝ^2^ subject to homogeneous Neumann (zero-flux) boundary conditions 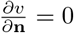 and 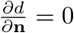 on the boundary ∂Ω where **n** is the outward normal vector. The system evolves from prescribed initial population distributions *v*(**x**, 0) = *v*_0_(**x**) and *d*(**x**, 0) = *d*_0_(**x**). All code is publicly available at https://github.com/shivmuthupandiyan/virus-dip-coevolution

## Results

### Numerical approach

To illustrate the model’s core dynamics, we solved the system numerically on a [-5, 5] *×* [-5, 5] domain with zero-flux boundary conditions. Unless otherwise specified, we used the following baseline parameters with units as given in Table 1: *µ* = 10^−3^, *η* = 10^−3^, *α* = 0.025, *β* = 0.025, *κ* = 10^−7^, *γ* = 0.3. The carrying capacity, *K*, was set to 10^8^ to give biologically reasonable population sizes [49]. Simulations were initialized with a virus population of 10^6^ and a DIP population of either 0 or 10^6^. The initial virus and (when present) DIP populations were distributed as Gaussians (standard deviation = 0.4) centered at (1, 1) and (1.4, 0), respectively. A population was considered extinct when its total size fell below 10^2^.

#### Baseline dynamics without interference

First, we consider a baseline scenario where virus infection produces non-interfering deletion variants as byproducts of viral replication; here the interference strength *κ* is set to zero (Fig 2a-c). These particles are not DIPS because they do not interfere with the virus. In this case, evolution of the virus population is driven solely by the intrinsic fitness landscape, and its mean phenotype moves toward the optimum at the origin. These mutants are generated from the virus at a rate *η* but cannot replicate on their own because the interference benefit *I*_*D*_ is zero. Consequently, the deletion mutant population is simply “dragged along” by the virus, remaining co-localized in phenotype space. Its population stabilizes when *de novo* generation (*ηv*) is balanced by decay (*γd*), yielding a population ratio of 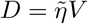. Without interference, there is no coevolutionary pressure, and the mutants act only as a minor sink on the virus population. This can map to a situation often observed during low multiplicity of infection (MOI) passages *in vitro*, where the few deletion mutants that are generated are not ever able to significantly establish themselves, causing minimal suppression of viral titers [50]. Here, the zero interference corresponds to passaging with sufficiently low MOI that the DIPs never coinfect the virus-infected cells, preventing any interference.

**Fig 2.**
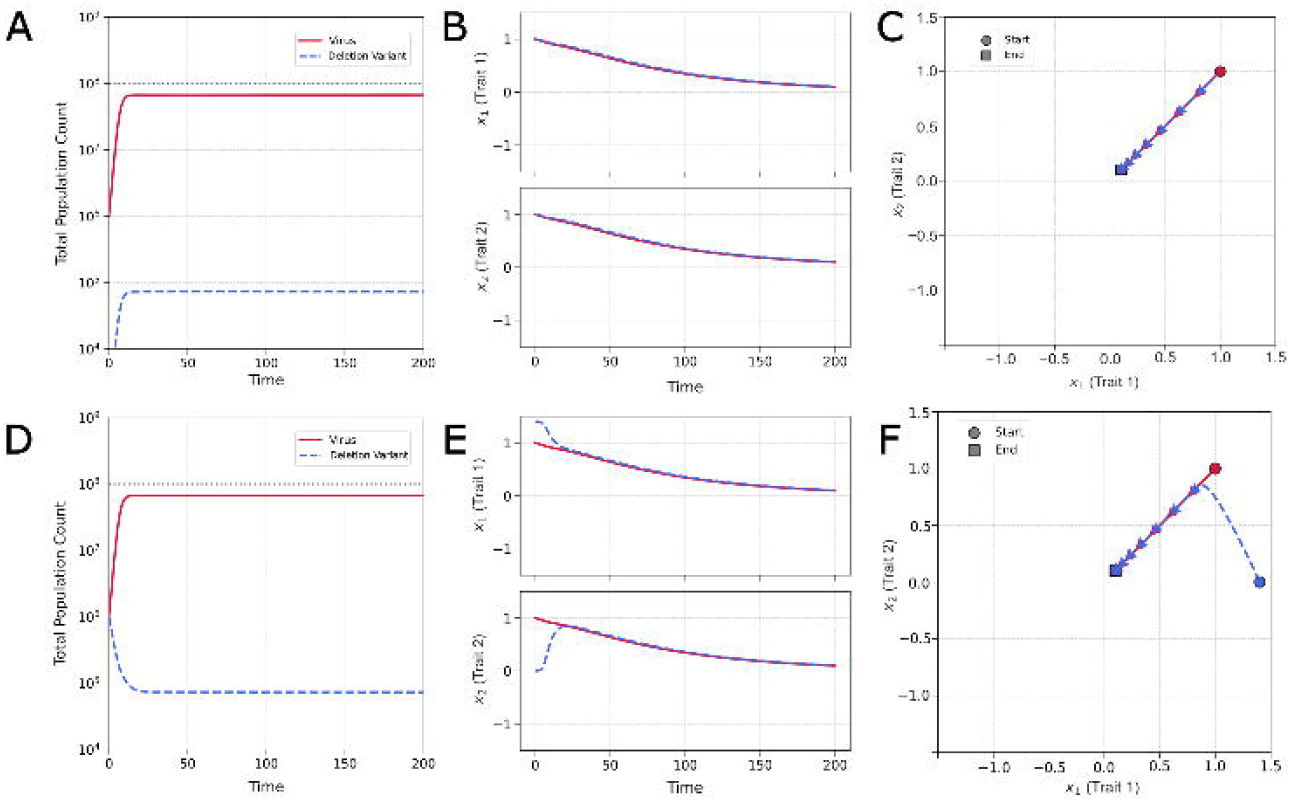
Virus population and evolutionary dynamics in the absence of interference by variants (*κ* = 0, *µ* = 10^−2^). **(A)** Baseline scenario with a deletion variant (blue dash) population emerging *de novo* from the virus (solid red) population. Total population sizes over time show viral dominance with deletion variant stabilizing at low levels. **(B)** Individual *x*_*1*_, *x*_*2*_ plots show both populations evolving toward the virus fitness optimum at the origin. **(C)** 2D phenotype trajectory demonstrates simple linear movement toward the optimum with virus and deletion variant populations remaining co-localized. **(D)** Robustness test with deletion variant populations initially phenotypically distant from the virus. Total population sizes converge to the same lowered level. **(E)** Deletion variant phenotypes converge to virus, while virus converges to origin. **(F)** 2D trajectory shows virus trajectory to origin and deletion variant trajectory to virus.

If the simulation begins with a non-interfering mutant population that is separate from the virus in phenotype space (Fig 2d-f), the initial population of mutants simply decays due to its inability to steal from the virus and replicate. It is steadily replaced by a *de novo* population that emerges co-localized with the virus. The phenotype centroid slowly shifts towards the virus centroid as more deletion mutants emerge from the virus, eventually driving the system back to the same steady state at the origin.

#### Interference drives coevolutionary chase

When interference is introduced (*κ >* 0), the dynamics become coevolutionary. DIPs can now replicate by parasitizing phenotypically similar viruses, creating a new selective pressure that disfavors viral phenotypes close to the DIPs. In a simulation where DIPs emerge *de novo* (Fig 3a-c), the virus experiences conflicting evolutionary forces: a directional selection towards the origin (the *α* term) and a selection away from the growing DIP population (the *κ* term). Without a fixed fitness peak (*α* = 0), the model would produce an “Escalatory Red Queen” dynamic, an endless arms race across a phenotype dimension. However, we assume that mutations that evade DIPs typically incur costs to intrinsic viral fitness, a phenomenon that has been observed in cases of antiviral and immune system escape [51]. The *α* term formalizes this constraint by establishing a global fitness optimum. This conflict results in a transient chase dynamic. As the virus nears the origin, the DIPs it generates become dense enough to repel it, causing the mean phenotype to overshoot the optimum before the *α* penalty pulls it back. While the trajectory of the population centroid is relatively simple, it arises from complex changes in the underlying distributions as they navigate this dynamic fitness landscape.

**Fig 3.**
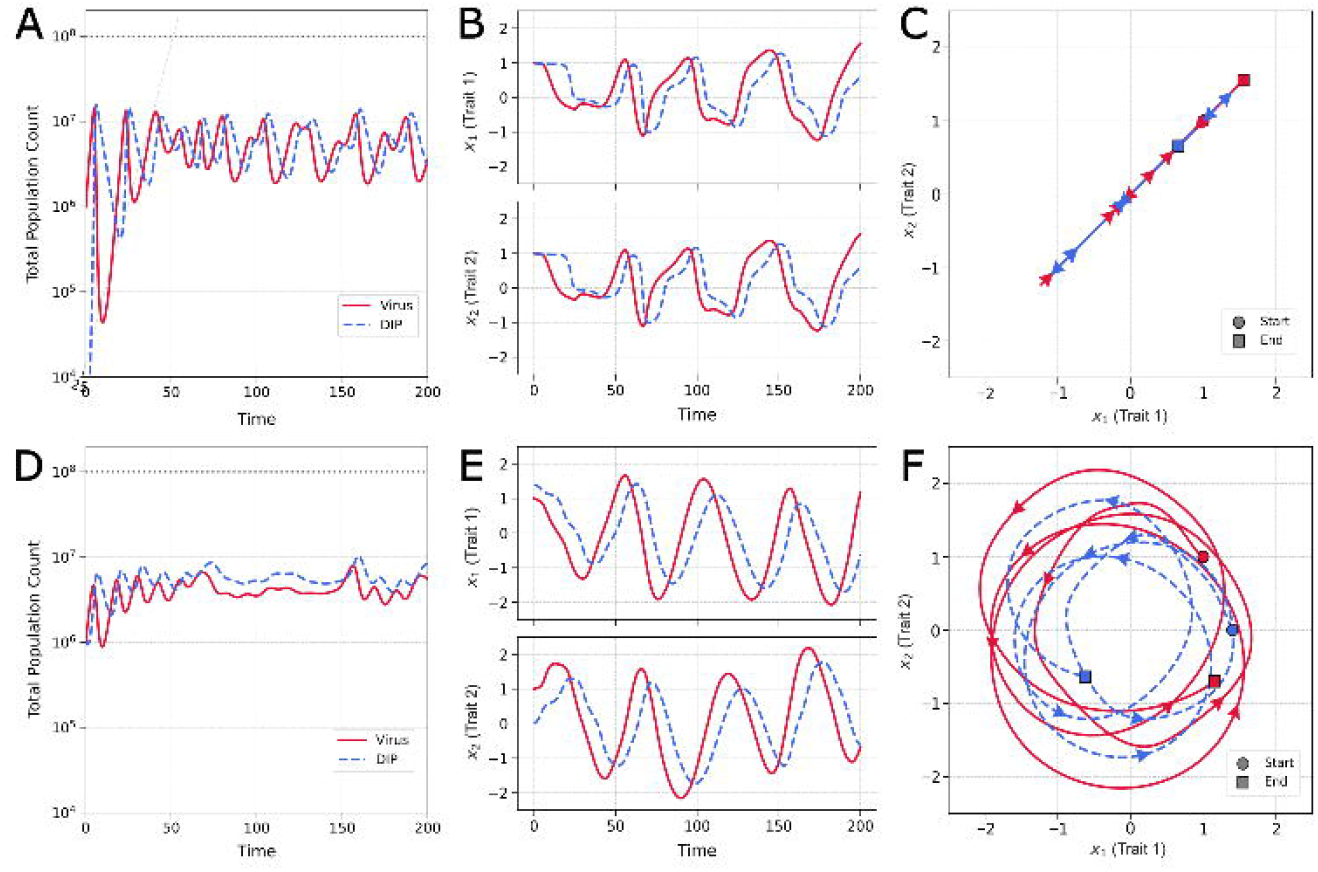
Population and evolutionary dynamics with virus-DIP interference (*κ >* 0, *µ* = 10^−2^). **(A)** Coevolutionary dynamics when DIPs (striped blue) emerge *de novo* from the virus (solid red) population. Oscillatory population dynamics demonstrate the von Magnus effect. **(B)** Phenotype trajectories show conflicting evolutionary pressures as the virus experiences a pull toward the fitness optimum (*α* term) and a push away from interfering DIPs (*κ* term), resulting in overshoot dynamics. Two-dimensional trajectory shows the oscillation along a line as the virus attempts to balance intrinsic fitness gains with interference avoidance. **(D).** Initial phenotypic separation induces a continuous coevolutionary pursuit where viruses evolve to escape DIP interference while DIPs track the moving viral target. Population oscillations reflect the dynamic strength of interference as phenotypic distance varies. **(E)** Large-amplitude oscillations in both phenotype dimensions demonstrate active coevolutionary chase. **(F)** Chase Red Queen trajectory in phenotype space.

Sustained coevolutionary chase dynamics emerge when the initial virus and DIP populations are phenotypically separated (Fig 3d-f). The initial offset breaks the linear trajectory seen previously, inducing a circular pursuit: viruses are selected to move away from the DIPs, while DIPs are selected to better track the viruses. The *α* term acts as a centripetal force, pulling the populations toward the origin and converting a linear arms race into a stable, cyclical pursuit. This dynamic, a chase across a multidimensional phenotype space, is the definition of a Chase Red Queen [38]. (See S1 Video)

#### Chase dynamics is reflected in both population size and phenotype

We observe the populations of DIP and virus oscillating over time (Fig 4a), a consequence of interference (von Magnus effect) that drives a predator-prey feedback observed in laboratory cell cultures of RNA viruses [5, 7, 8] as well as in mice [6]. High levels of DIPs suppress the virus; this depletes the resources available for DIP replication, causing the DIP population to fall; the virus population is then able to recover, restarting the cycle.

**Fig 4.**
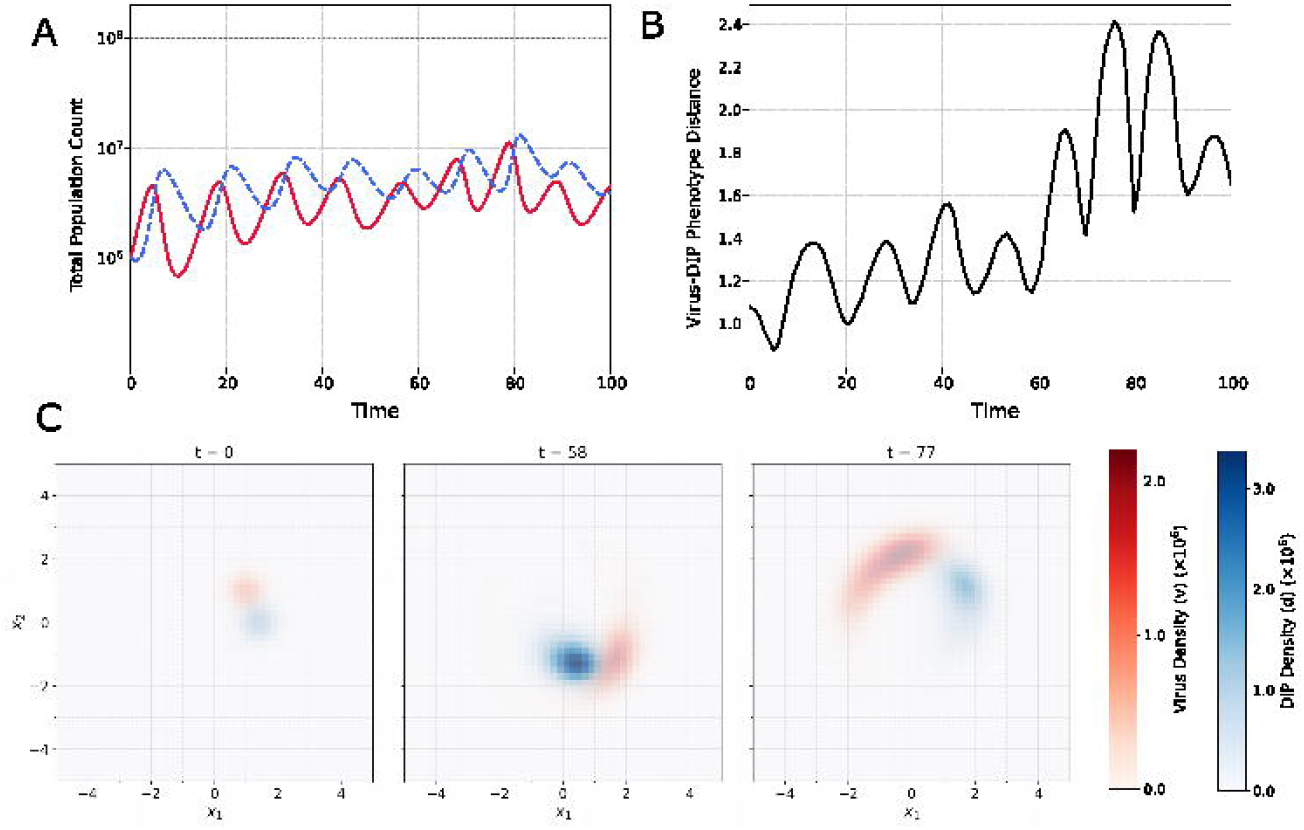
Coevolutionary chase is associated with oscillations in populations and phenotype distances. **(A)** Oscillations in virus and DIP populations over time exhibit predatory-prey population cycles. **(B)** Distance between mean phenotypes oscillates over time, creating periods of DIP containment and viral escape **(C)** Heatmaps of DIP and virus densities in phenotype space show initial conditions (t=0), minimum distance (t=58), and maximum distance (T=77). These illustrate the full viral distribution, rather than a single point mean.

Our model extends this dynamic by allowing the strength of virus–DIP interference to depend both on phenotypic distance and on the population sizes of *V* and *D* via the interference terms of Eqs (2) and (1), respectively, with phenotypic distance itself oscillating as the virus escapes and the DIP gives chase. To quantify the chase dynamics, we track the distance between the centroids of the virus and DIP populations (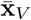and 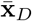) over time (Fig 4b). The chase dynamics, continuous cycles of viral escape and DIP pursuit, directly influence population sizes. These cycles are not always synchronized: viral recovery can occur either after a DIP crash or by creating sufficient phenotypic distance to weaken interference, linking population cycles to evolutionary dynamics.

Snapshots of the population densities reveal the coevolutionary chase in detail (Fig 4c). At any given time, both the virus and DIP populations exist not as single points but as diffuse clouds in phenotype space. This distribution is analogous to a viral quasispecies, where a central dominant phenotype is surrounded by a spectrum of mutants. The heatmaps illustrate the continuous pursuit, with the DIP cloud tracking the virus cloud as it moves to evade interference (See S2 Video).

The sustained chase dynamic shown in Figs 3 and 4 represents one of several possible outcomes. Depending on the model parameters, the coevolutionary interaction can instead lead to DIP extinction, virus-DIP coextinction, or a stable coexistence with no oscillations. There is no region for virus extinction alone, as DIPs cannot replicate in the absence of virus, so virus extinction entails DIP extinction. The next section explores how key parameters determine which of these regimes will emerge.

### Sensitivity analysis

We performed a global sensitivity analysis using Latin Hypercube Sampling (LHS) (n = 10,000) to explore how variation in six key parameters shapes the prevalence of distinct coevolutionary regimes. For each parameter, outcomes were classified as Chase, Coexistence, DIP Extinction, or Coextinction. We swept parameter ranges: *κ* : (10^−1^, 10^2^), *η* : (5 *×* 10^−4^, 5 *×* 10^−1^), *α* : (3 *×* 10^−3^, 3), *β* : (3 *×* 10^−3^, 3), *γ* : (0, 0.9), *µ* : (10^−4^, 10^−2^), all on log scales except for *γ*. Populations were initialized at 10^−2^ particles for both virus and DIP, with carrying capacity at 1. Population extinction thresholds were set at 10^−6^, with simulations classified as ‘DIP Extinction’ when DIP populations fell below this value and as ‘Coextinction’ when viral populations fell below this value. Dynamics were classified as ‘Chase’ when the variation in Euclidean distance between DIP and virus distribution centroids exceeded 0.1, and as ‘Coexistence’ otherwise.

Parameters were selected to avoid trivial extinction and encompass biologically plausible dynamics. The resulting stacked proportion plots (Figs 5a-5f) reveal how the likelihood of each regime changes as individual parameters vary, while accounting for uncertainty in all other parameters.

**Fig 5.**
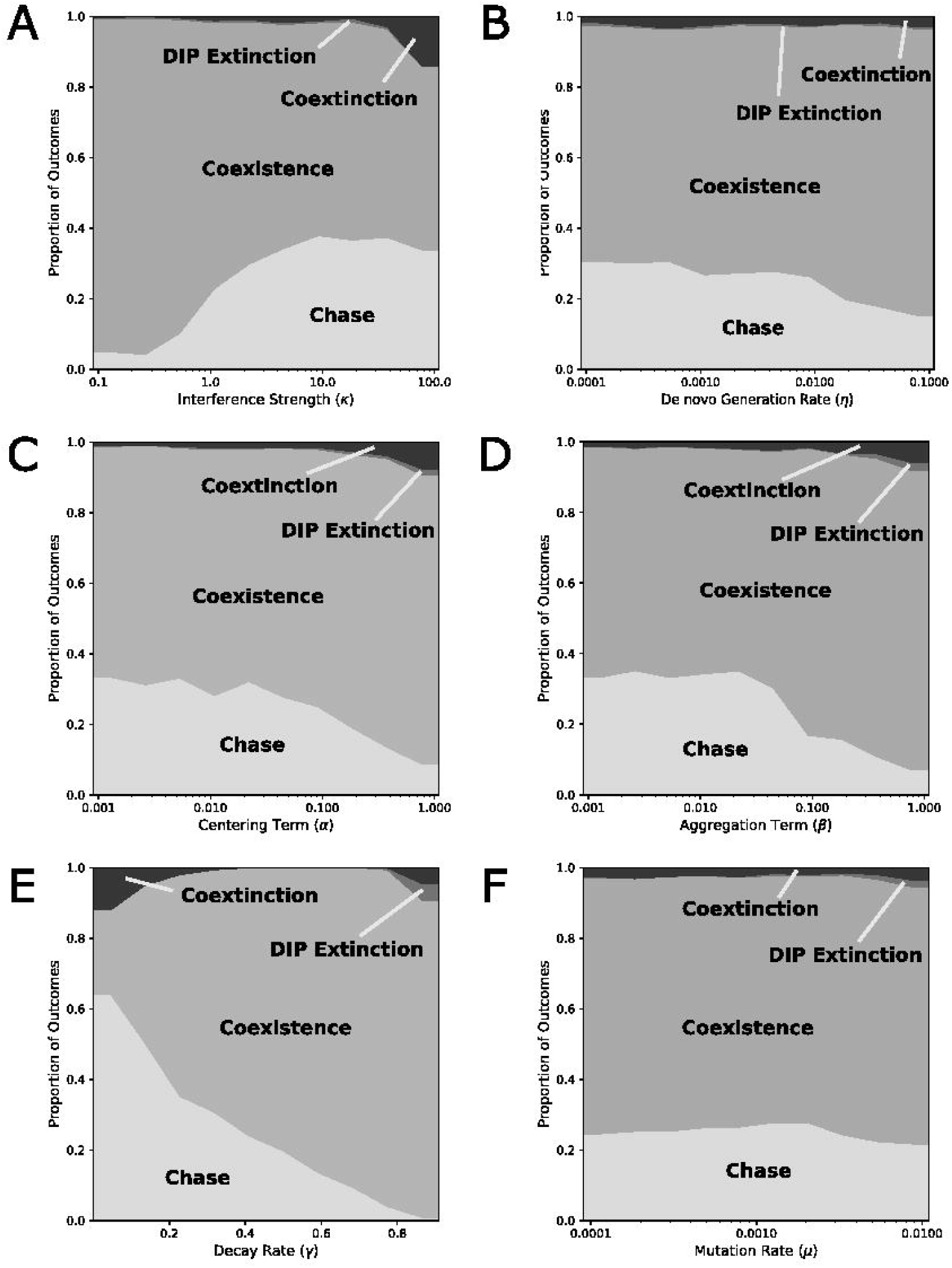
Sensitivity analysis of the nondimensionalized model. (A) Interference strength (*κ*). Increasing the interference strengths of DIPs burdens the virus, shifting populations from coexistence towards chase regimes, until interference is so strong that viruses start to go extinct. **(B) *De novo* generation** (*η*). Larger *de novo* emergence of DIPs colocalized with virus slightly favor coexistence over chase outcomes. **(C) Centering Term** (*α*). Drawing the virus closer to its maximum fitness (origin), suppresses its ability to escape, favoring coexistence and extinction outcomes. **Aggregation Term** (*β*). Increasing the aggregation discourages virus phenotypic diversification, as mutations that drive diversification are disfavored. Effects mirror those of the centering term. **(E) Decay Rate** (*γ*). At very low decay rates, virus and DIP growth approach their carrying capacity because there is very little washout, leading to chase outcomes and coextinction. As the decay rate increases, variation is selected against more strongly, preventing a chase from occurring, until very high decay rates, when DIP extinction and coextinction rise again. **(F) Mutation rate** (*µ*). Mutation rates have a negligible effect in the examined parameter range, as the selection is much stronger in the mutation-selection balance.

Several robust patterns emerge. Interference strength (*κ*, Fig 5a) shows a clear progression: when *κ* is low, nearly all simulations result in coexistence, as interference is too weak to generate selective pressure. At intermediate *κ*, chase dynamics become possible (~ 20% of outcomes), but excessively strong interference pushes the system toward coextinction. The *de novo* DIP generation rate (*η*, Fig 5b) has an equally strong effect: at low *η*, DIPs fail to persist and chase is common, but higher values reduce the opportunity for phenotypic separation by continuously regenerating DIPs near the virus, enforcing coexistence.

The phenotypic sensitivity parameters (*α*, Fig 5c; *β*, Fig 5d) both display similar trade-offs. Low values favor chase by allowing viruses to maintain phenotypic separation, while higher values suppress coevolution and shift the system toward coexistence and extinction outcomes. This symmetry suggests that both parameters act as modulators of how tightly viral and DIP dynamics are coupled.

The washout rate (*γ*, Fig 5e) exhibits the most complex effects. At low values, chase is highly prevalent (nearly 50%) alongside elevated extinction rates, reflecting that limited washout disproportionately benefits DIPs and forces the virus into rapid adaptation or collapse. By analogy with a bioreactor, allowing the washout rate to approach zero means the influx of fresh cells for infection or coinfection approaches zero, and the system approaches a closed system, where carrying capacity is used and the logistic factor *L* approaches zero. As *γ* increases, extinction decreases and coexistence grows more common, indicating that faster removal weakens DIP pressure. However, at the highest values, strong washout suppresses both populations and again increases extinction. This reveals that *γ* tunes the relative balance of viral escape and DIP persistence in a strongly nonlinear manner.

Mutation rate (*µ*, Fig 5f) has comparatively weaker effects: chase maintains a consistent prevalence of ~ 25%, while coexistence dominates (~ 70%). DIP extinction and coextinction occur only rarely in this range. The relatively low frequency of chase here reflects the chosen parameter bounds; other ranges could yield a different balance.

Overall, the LHS sensitivity analysis shows that chase dynamics are most common under intermediate interference strength, low-to-moderate DIP generation, and low washout. Coexistence dominates large regions of parameter space, especially when interference is weak, mutation rate is moderate, or washout is strong. Extinction outcomes emerge primarily at the extremes of parameter values. Together, these results highlight that coevolutionary chase requires a narrow balance of forces, while Coexistence and extinction represent more robust attractors of the virus–DIP system.

### Cyclical escape

Our model’s coevolutionary chase dynamics provide a mechanistic explanation for classic experimental observations of cyclical viral resistance. In time-shift experiments, DePolo et al. found that VSV is most resistant to DIPs from the recent past, yet remains susceptible to DIPs from the distant past and future [31]. This pattern of transient resistance argues against a simple Escalatory Red Queen arms race, where resistance to past DIPs would be expected to increase monotonically. Instead, it is a hallmark of a Chase Red Queen dynamic, which our model conceptualizes as a cyclical pursuit within a constrained phenotype space. Because the evolutionary trajectory is orbital, a virus population at a given time can be phenotypically closer to DIPs from the distant past or future than to the DIPs from the recent past it has just evolved to escape.

To test this in our model, we first generated a heatmap showing how viral resistance to DIP interference varied over time (Fig 6a). The diagonal stripes reveal alternating cycles of high and low resistance, indicating ongoing coevolution between the virus and DIP populations. We then fixed one DIP population (from time T = 77) and measured its interference against viruses from surrounding passages (Fig 6b). The virus was least resistant to DIPs from its own time, more resistant to those from the recent past—having just evolved to escape them—and less resistant again to those from the distant past or future. Finally, to compare these predictions with experimental data, we devised a time-shift sampling approach to mimic the DePolo serial transfer experiments (Fig 6c). The resulting pattern matched the transient cycles of resistance observed in the original study.

**Fig 6.**
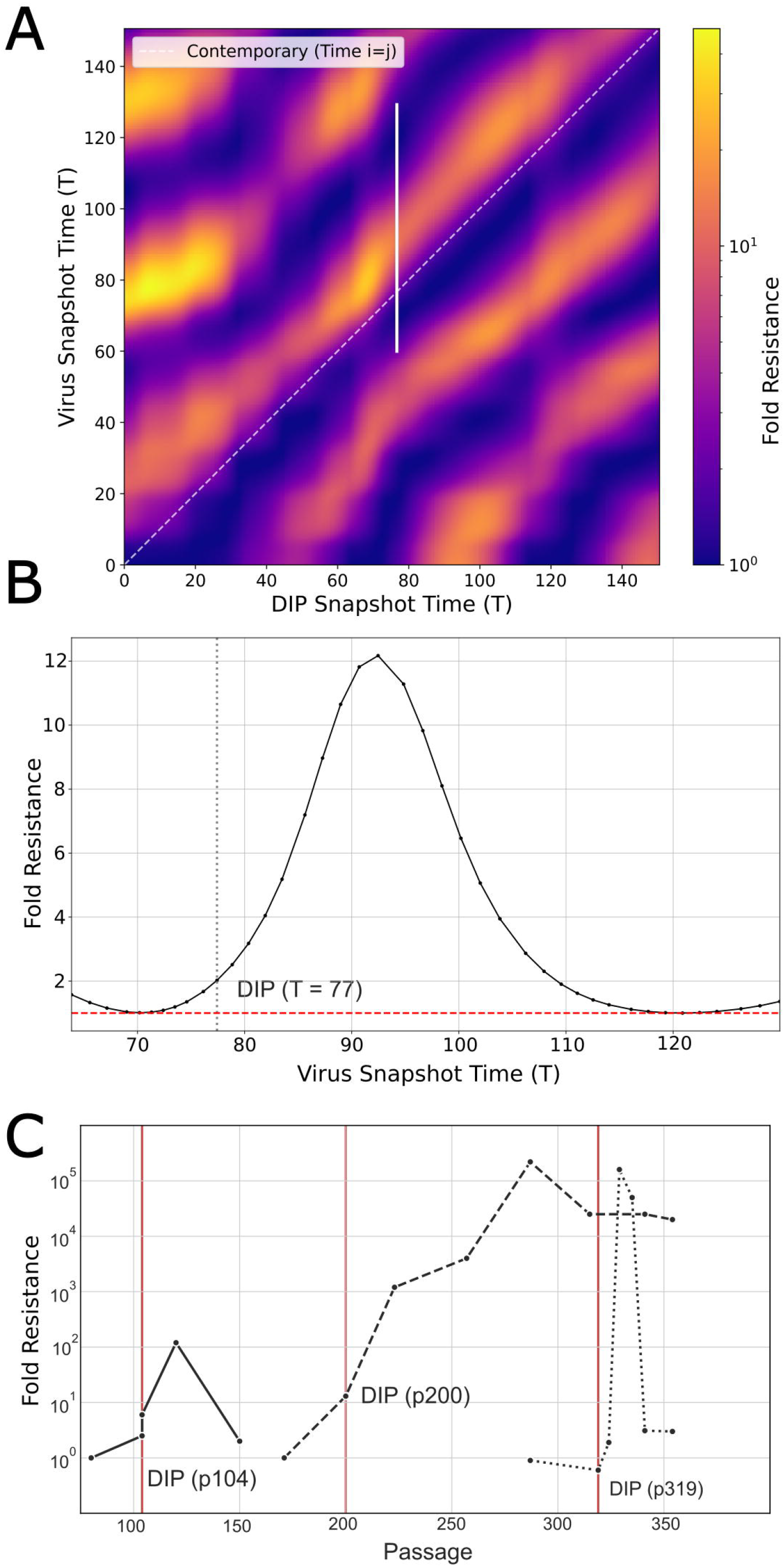
Cross passage viral resistance. **(A)** Resistance of any given virus passage (y-axis) against any given DIP passage (x-axis), measured as fold reductions in interference. The white diagonal represents the resistance against contemporary passage (DIP Snapshot Time T_i_ = Virus Snapshot Time T_j_). The diagonal stripes represent a coevolutionary chase where viruses are most resistant to DIPs from the recent past, before losing their resistance. The vertical white line shows the subset of resistances depicted in B. **(B)** Example viral resistance curve against a specific DIP (T=77), showing a transient increase in resistance for several passages, before reverting to the baseline resistance (red dashed line). The domain corresponds to the vertical white line in A. **(C)** Replotting of data from DePolo et al. (1987), showing viral resistance to DIPs cycling over several passages.

### Evolutionary conflict

Our model also provides a framework to examine “evolutionary conflict,” an emerging concept in the design of therapeutic interfering particles (TIPs). In a multi-scale model of HIV, Rast et al. proposed that TIPs could create an evolutionary trap: any viral mutation that evades interference also incurs a significant fitness cost, blocking the evolution of resistance [27]. In our model, this constraint is captured by the *α* parameter, which penalizes deviations from the fitness optimum at the origin.

This raises a key question: under what conditions does the selective pressure to escape TIPs (*κ*) outweigh the fitness cost of deviating from the optimum (*α*)? To probe this trade-off, we examined virus population sizes and virus-DIP phenotypic distances across a range of *κ* and *α* values (Fig 7). As expected, virus populations decline with increasing interference by DIPs, largely independent of *α* (Fig 7a). However, the average phenotypic distance between virus and DIP centroids reveals a clear threshold behavior (Fig 7b). When the fitness penalty for deviation is strong (e.g., *α >* 0.5), the virus remains “trapped” at the origin regardless of the interference strength. Any mutation away from the optimum is so costly that the virus cannot stably evolve any DIP resistance. The distance remains constant even as the interference strength increases (Fig 7b), and the virus population rapidly decreases (Fig 7a). Below this *α* threshold, however, a sufficiently high interference cost (*κ*) forces the virus to abandon the optimum and engage in a coevolutionary chase (Fig 7b). This escape has direct consequences for viral control. In the “trapped” regime (high *α*), increasing *κ* effectively suppresses the viral population. In the “escape” regime (low *α*), the virus population is less affected by *κ* because it can simply evolve away from the interference with minimal loss of fitness.

**Fig 7.**
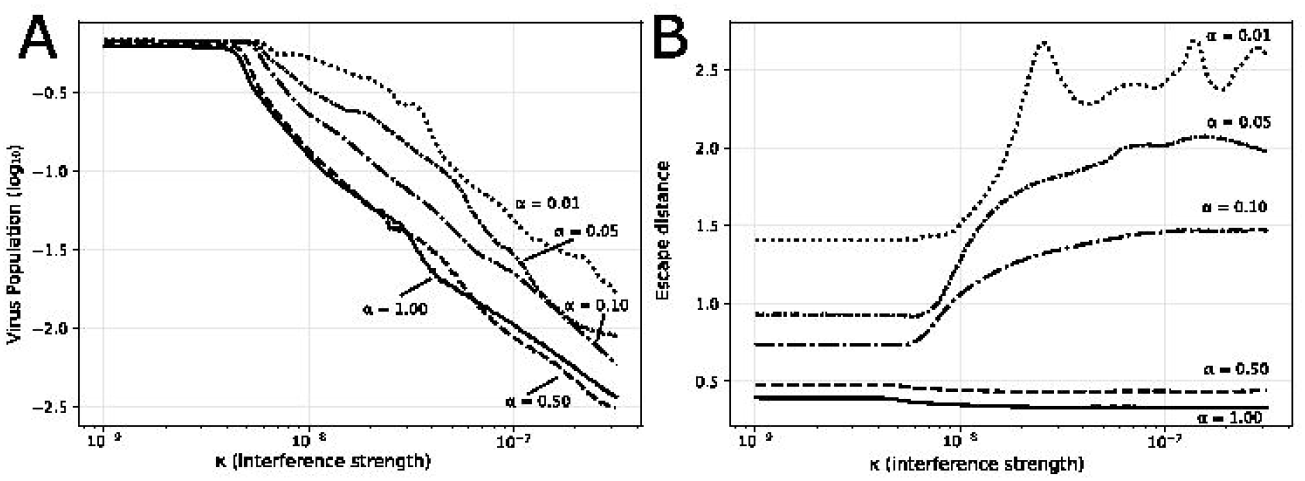
Tradeoff Between Viral Fitness and Escape. **(A)** Final virus population size versus interference strength (*κ*) for different fitness penalty parameters (*α*) from (dotted) to 1 (solid). Higher *α* values causes the decline to be more pronounced, as the virus is less able to escape as the interference becomes stronger. **(B)** Extent of viral escape (mean distance between virus and DIP centroids) versus interference strength. High *α* values maintain viruses near the optimum regardless of *κ* (flat lines), representing successful evolutionary containment. Low *α* values show increasing escape with interference strength, demonstrating that evolutionary conflict strategies succeed only when escape mutations incur prohibitive fitness costs (high *α*), but fail when lower-cost escape pathways exist (low *α*).

Our model therefore suggests that the success of an evolutionary conflict depends critically on the fitness cost of escape mutations. The containment described by Rast et al. is effective because its modeled escape pathways, such as reduced capsid production, incur a large intrinsic fitness cost to the virus. This scenario is represented by the high-*α* regime in our model, where the virus is successfully contained.

Our model demonstrates, however, that such containment may fail if the virus can find escape routes with a lower fitness penalty (the low-*α* regime). This scenario is exemplified by VSV, which evolved resistance to defective interfering particles through mutations in its polymerase complex. These mutations selectively reduced the polymerase’s affinity for interfering templates while preserving its function on the viral genome, allowing escape with a minimal fitness cost [32]. This example highlights a fundamental asymmetry in the evolutionary arms race: the virus only needs to find one viable escape route within a vast, high-dimensional landscape of possibilities, whereas the TIP must be robust to all of them. Therefore, the long-term success of a TIP-based therapy likely depends on targeting a viral function so essential that any resistance mutation incurs a prohibitive fitness cost.

## Discussion

We developed a mathematical model to investigate the coevolutionary dynamics between a virus and its defective interfering particles. By integrating phenotype-dependent interference, mutation, and intrinsic fitness landscapes, our framework demonstrates how a limited set of virus-DIP interactions can generate a wide spectrum of complex outcomes. The model successfully reproduces key phenomena observed in experimental systems, including oscillating population dynamics (stemming from the von Magnus effect) and sustained coevolutionary chases (Red Queen). The emergence of these behaviors from the model’s core assumptions provides a powerful tool for exploring the evolutionary principles that govern viral resistance to interfering particle therapies.

Our model’s explanatory power is limited by its core structural assumptions. We defined viral interaction using a Gaussian interference kernel and implemented quadratic fitness terms (*α, β*). While these are mathematical abstractions, the model’s behavior is not wholly dependent on them; substituting alternative kernels (e.g., top-hat, exponential) produced qualitatively similar chase dynamics (Fig S5), suggesting the results are robust.

Further simplifications relate to the model’s scope. We represent the vast genetic and phenotypic landscape of a virus in only two dimensions. In reality, dimensionality is likely much higher when taking into account complex interference mechanisms, like the activation by DIPs of host immune defenses [12, 52] or the creation of novel chimeric proteins and functions as a byproduct of DIP formation [28, 53–58]. Here too, our core findings hold under limited testing: simulations in three trait dimensions showed similar behaviors to two dimensions, though computational costs prohibited a thorough analysis (Fig S4). We use a deterministic framework that neglects the randomness of evolution, but assume that in large populations, the stochastic effects are less prominent. Our modeling of distance using simple Euclidean measures between population centroids omits information about the shape of the distributions. However, we find similar trends when using a more comprehensive distance measure (Fig S3).

Applying our modeling approach to a specific virus system will require translating its abstract phenotype space into one defined by concrete, measurable biological traits. For instance, the trait axes may represent features such as the viral replicase or capsid protein binding affinities to the virus and DIP genomes. Such a specialization would necessitate modifying the model’s core assumptions. Mutation, for example, may no longer be isotropic, as certain traits may be inherently more sensitive to mutation than others. Furthermore, representing virus and DIP traits in separate phenotype spaces to capture virus or DIP specific parameters (e.g., DIP genome length) would demand a redesign of the interaction kernels. While these adaptations would enhance biological realism, they would do so at the cost of the model’s generalizability. Therefore, the abstractions of the current model are best viewed as a foundational framework, designed to be adapted and specialized for future, system-specific investigations.

For instance, while our model qualitatively reproduces observed transient resistance patterns, there are quantitative differences from the experimental data (Fig 6). The modeled resistance changes are on a linear scale, not the orders-of-magnitude changes experimentally observed. Additionally, the model neglects the constant accumulation of mutations observed in regions like the N gene and 5’ termini, which prevent the virus from returning exactly to previous phenotypes [31]. These discrepancies are expected consequences of model simplifications, such as the two-dimensional phenotype space, which constrains evolutionary trajectories. Within our framework, the drift and orders-of-magnitude resistance could be represented as a “corkscrew” trajectory through higher-dimensional space, where cyclical pursuit in some dimensions is coupled to directional drift in others. Such an extension could be implemented by expanding the phenotype space or allowing the quadratic fitness optimum in the *α* term to drift slowly over time. While promising, these ideas lie beyond our present goal of isolating the core chase dynamics. The model’s value here is not in precise numerical replication, but in its ability to predict how the qualitative chase dynamic responds to changes in biological parameters.

A clear path forward is to connect the model’s abstract parameters to experimentally measurable quantities. Parameters like mutation rate (*µ*) and decay rate (*γ*) can often be measured directly or manipulated in laboratory systems [59, 60]. Others, such as the interference strength (*κ*), would require more sophisticated experimental designs.

Experimental observations align well with our model’s predicted trends. In particular, the shift from coexistence to chase dynamics as interference strength increases (Fig 5A) is supported by laboratory evolution studies. Pelz et al. tracked DIPs of influenza A in long-term culture and found that the variants that eventually dominated—and drove oscillatory population dynamics—were those with greater interfering efficacy [61]. This selection for highly potent DIPs in a dynamic context mirrors the “chase” regime of the model, which arises only when *κ* is high enough to overcome coexistence constraints.

A compelling experimental test of our model’s predictions would be to compare the coevolutionary dynamics of different virus-DIP systems. For example, bioreactor systems with higher outflow rates should, according to our model, exhibit more stationary evolutionary dynamics than those with lower turnover rates (Fig 5e). Such comparative studies would provide a robust validation of the core principles identified in our theoretical framework.

The sensitivity analysis reveals that successful TIP design may require navigating several trade-offs rather than simply maximizing any single parameter. The most robust therapeutic outcomes occur in parameter ranges with high interference and high costs to evolutionary escape (Fig 7). These lead to the ideal scenarios of: (1) complete virus extinction or (2) established stable coexistence with ongoing but manageable viral suppression. The intermediate chase regime, while evolutionarily interesting, may be less desirable therapeutically due to its oscillatory viral loads and potential for eventual viral escape. Critically, these results suggest that TIP failure is most likely to occur through low costs to escape (e.g., targeting non-conserved viral functions in high-mutation environments) or insufficient interference strength relative to viral fitness advantages.

In summary, this work presents a framework for understanding virus-DIP coevolution not as a simple arms race, but as a complex interplay of forces that can lead to diverse outcomes. The model’s primary contribution is to translate abstract evolutionary pressures into distinct, predictable regimes. By identifying the conditions that lead to stable suppression, viral escape, or sustained coevolutionary chases, our model provides a conceptual foundation for generating testable hypotheses. Ultimately, this approach may help guide the rational design of therapeutic strategies aimed at anticipating and steering viral evolution.

## Acknowledgments

We gratefully acknowledge support by the National Science Foundation (NSF) under grants DMS-2151959, MCB-2029281, and CBET-2030750; the National Institutes of Health (NIH) under awards OT2OD030524 and R01DK133605; and institutional support from the University of Wisconsin–Madison, including the Wisconsin Institute for Discovery, the Office of the Vice Chancellor for Research, and the Department of Chemical and Biological Engineering.

## Supporting information

**S1 Video. Video visualization of simulation in Fig 3b**.

**S2 Video. Video visualization of simulation in Fig 4**.

**S3 Figure. Earth mover’s distance**. A simple Euclidean distance can be misleading, as it overlooks changes in the shape and spread of the population distributions. We therefore also compute the Earth Mover’s Distance (EMD), which measures the total “work” required to transform the full virus distribution into the DIP distribution. The EMD provides a more comprehensive measure of their dissimilarity and also exhibits clear oscillations, validating that the chase dynamic involves the entire quasispecies, not just the movement of its mean.

**S4 Figure. Higher dimensional modeling**. We extend the model to function in three dimensional space. We initialized the models with only a small perturbation in the 3rd dimension. From that, we saw that the perturbation expanded to the higher dimension, rather than any collapse or full escape, suggesting that the chase modeling can be stability extended to higher dimensionality.

**S5 Figure. Alternative interference kernels**. The main text model uses a Gaussian interference kernel 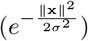, to capture the intuition that more similar DIPs will better be able to interfere with the virus. However, we relax this assumption and test several other kernels. These include exponential (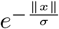, slower decay) (d-f), rational (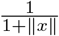, much slower decay) (g-i), and step function (1 if ∥*x*∥*< σ*, else 0) (j-l). For the most part, these different kernels yield distinct but not disparate trajectories, suggesting that the model is robust to the choice of interference kernel.

