## Supplementary figures and images for "Coevolutionary dynamics of viruses and their defective interfering particles"

### Supplemental Figure3

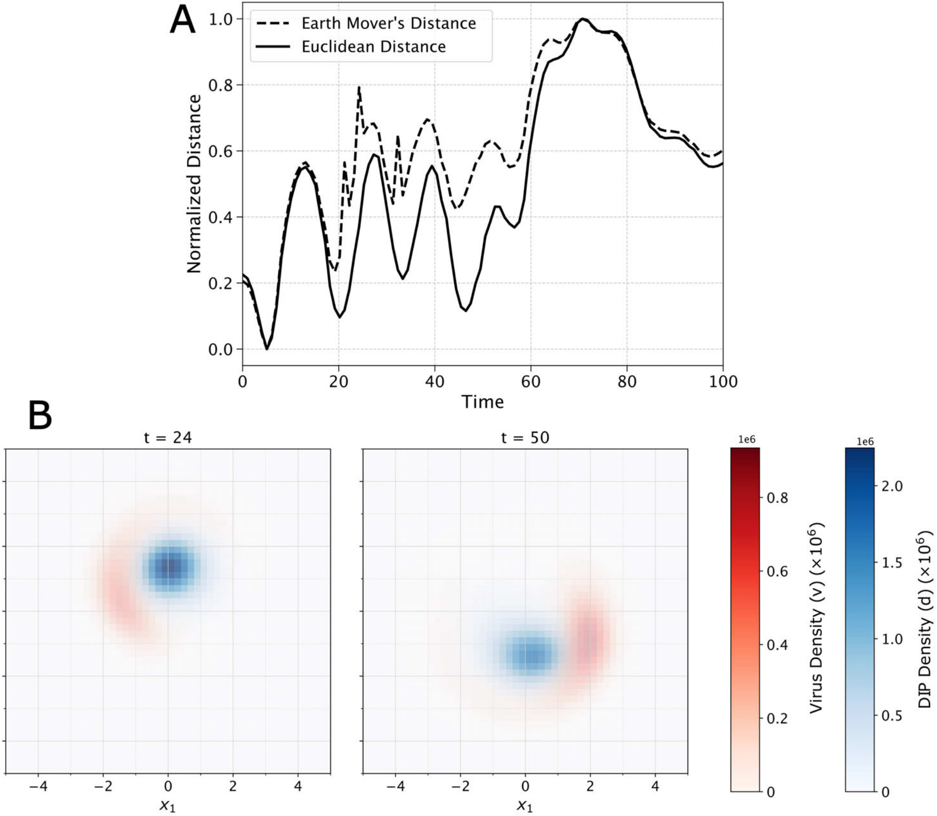

### Supplemental Figure4

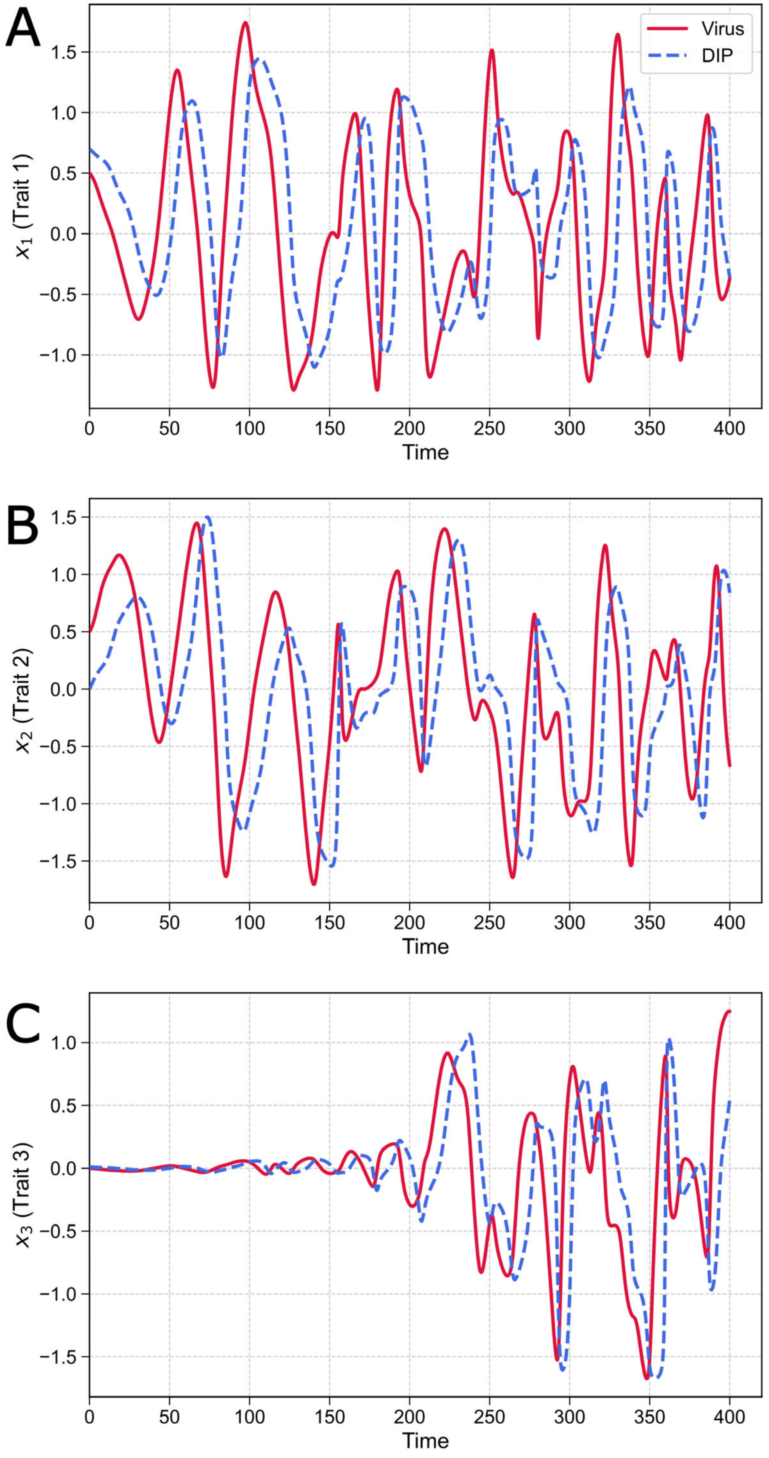

### Supplemental Figure5

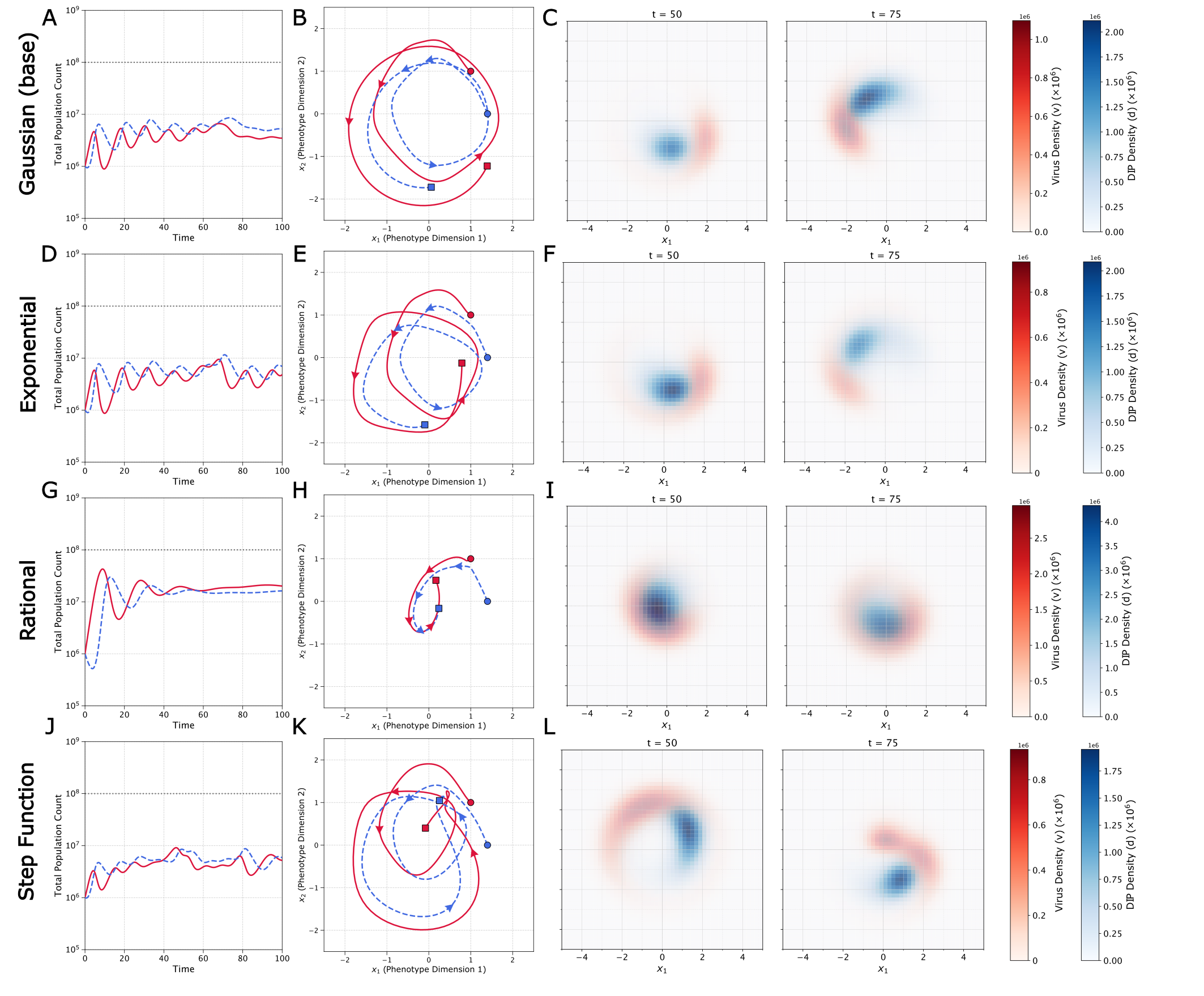
